# Evaluation of FRET X for Single-Molecule Protein Fingerprinting

**DOI:** 10.1101/2021.06.30.450512

**Authors:** Carlos de Lannoy, Mike Filius, Raman van Wee, Chirlmin Joo, Dick de Ridder

**Affiliations:** Bioinformatics Group, Wageningen University, Droevendaalsesteeg 1, 6708PB, Wageningen, The Netherlands; Department of BioNanoScience, Kavli Institute of Nanoscience, Delft University of Technology, van der Maasweg 9, 2629HZ Delft, The Netherlands

## Abstract

Single-molecule protein identification is a novel, as of yet unrealized concept with potentially groundbreaking applications in biological research. We propose a method called FRET X (Förster Resonance Energy Transfer via DNA eXchange) fingerprinting, in which the FRET efficiency is read out between exchangeable dyes on protein-bound DNA docking strands, and accumulated FRET efficiency values constitute the fingerprint for a protein. To evaluate the feasibility of this approach, we simulated fingerprints for hundreds of proteins using a coarse-grained lattice model and experimentally demonstrated FRET X fingerprinting on a system of model peptides. Measured fingerprints are in agreement with our simulations, corroborating the validity of our modeling approach. In a simulated complex mixture of >300 human proteins of which only cysteines, lysines and arginines were labeled, a support vector machine was able to identify constituents with 95% accuracy. We anticipate that our FRET X fingerprinting approach will form the basis of an analysis tool for targeted proteomics.

## Introduction

Proteins come in a wide variety of shapes, sizes and forms. Each is attuned to fulfill one or more of the many functions that are essential to living cells, including the catalysis of metabolic reactions, replication of genetic information, provision of structural support, transport of molecules and many more. To fully understand the biological processes taking place in a cell, it is critical to identify and quantify constituents of its proteome at any given time during the cell cycle.

Mass spectrometry (MS) is currently the gold standard for protein identification and quantification. Over the past decades, MS techniques have improved tremendously in terms of accuracy and dynamic range; however, detecting and distinguishing all proteins in complex samples remains challenging. Many biologically and clinically relevant proteins such as signaling molecules and disease biomarkers occur in such low abundance that they remain undetectable by MS.^1^ Moreover, the proteome complexity increases through alternative splicing or posttranslational modifications, as a single gene can produce dozens of distinct protein varieties, referred to as proteoforms.^2^ Not all of these proteoforms canbe distinguished by current approaches. As such, there is considerable incentive for the development of new protein sequencing methods that operate at the single-molecule level.^3,4^

Single-molecule techniques have boosted DNA sequencing, allowing for the identification of individual nucleic acid molecules, and are now routinely used for genome and transcriptome mapping of single cells.^5^ However, the search for single-molecule protein sequencing techniques is not trivial due to the high complexity of protein molecules compared to DNA molecules. For example, the DNA code consists of only four nucleotides whereas there are twenty different amino acids for proteins. Furthermore, low abundant DNA molecules can be enzymatically amplified outside the cell whereas such an enzyme is absent for proteins.

Novel single-molecule protein analysis methods have been proposed to circumvent this additional complexity. Importantly, only a subset of the theoretically possible combinations of polypeptide chains occurs in nature, and a fraction of that subset is of importance in a given research setting. Therefore, proteins may be identified by reading out a signature of incomplete information, which is then compared to a database of relevant signatures. We refer to this approach as protein fingerprinting, and to said protein signatures as protein fingerprints. It has been shown that sufficiently distinct protein fingerprints only require the read-out of a small subset of residue types.^6-8^ If cysteine and lysine residues were orthogonally labeled and read out sequentially, our simulations indicated that the majority of human proteins were uniquely identifiable.

Several novel protein fingerprinting methods based on the readout of a subset of residue types have recently been demonstrated, most of which require linearization of the polypeptide chain to allow for the determination of the residue order.^9,10^ This linearization can be achieved by translocating the polypeptide chain through a nanopore^4^ or by using a fluorescently labeled motor protein^8^ to read out the modified residues required for fingerprinting. Alternatively, the protein fingerprint can be obtained by labeling certain amino acids and determining their location through several Edman degradation cycles^11^. Although full length proteins are difficult to analyze due to the limited number of Edman cycles that can be performed, its utility for analyzing shorter peptides has been shown in a proof of concept. All these approaches have in common that they probe each protein only once, while the accuracy would increase if the same molecule could be measured multiple times.

In this study, we present a novel protein fingerprinting method that builds further on the concept of residue-specific labeling of selected amino acids and obtains a protein fingerprint by determining the location of amino acids in the 3D structure of a protein. As the size of most proteins lies in the low nanometer range, our protein fingerprinting approach requires a technique that can determine the location of residues with sub-nanometer resolution. Single-molecule FRET is well suited for this and comes with the benefit that several thousands of molecules can be imaged at the same time, if full length proteins can be immobilized in a microfluidic chamber.^12^ Here we verify the feasibility of a single-molecule FRET-based protein fingerprinting method. We first demonstrate that experimentally obtained fingerprints for four model peptides are distinct and are reproduced by our simulation method. Then we show that simulated fingerprints of 313 human proteome constituents can be identified with 95% accuracy. If mislabeling of residues is assumed to occur, this accuracy decreases to 91%. This supports the notion that FRET fingerprinting allows for the reliable identification of proteins in complex mixtures.

## Approach

### FRETX for protein fingerprinting

To realize protein fingerprinting using single-molecule FRET, a resolution sufficient to determine the location of multiple amino acids in the protein structure is required. However, single-molecule FRET analysis is limited to just one or two FRET pairs in a single measurement.^13,14^ Recently, our group developed a concept to allow for the detection of multiple FRET pairs in a single nanoscopic object. Our technique, FRET X (FRET via DNA eXchange), employs transient hybridization of DNA strands labeled with a fluorophore to temporally separate FRET events that originate from different FRET pairs. We have shown that FRET X can resolve the distance between multiple FRET pairs with sub-nanometer accuracy.^15^ Here, we apply FRET X for protein fingerprinting. By detecting target amino acids one by one, FRET X produces a unique fingerprint, allowing identification of the protein from a reference database.

**Figure 1** illustrates the workflow for protein fingerprinting using FRET X. A subset of amino acids of a protein of interest is labeled with orthogonal DNA sequences, which serve as docking strands for their complementary imager strands (**Figure 1A**). One of the termini is labeled with a unique DNA sequence, which functions as a reference point and facilitates immobilization of the full-length protein to a microfluidic chip. To obtain a FRET fingerprint for one of the amino acids, fluorescently labeled imager strands for the terminal reference sequence and for the particular amino acid (e.g. Cysteine, **Figure 1B**) are added. The imager strands for the reference point are labeled with an acceptor fluorophore, while those for the cysteines carry a donor. FRET can occur only when both imager strands are simultaneously bound. The transient and repetitive binding of imager strands reports on the relative location of a residue to the reference point. Furthermore, since the pool of fluorophores is continuously replenished, the effect of photobleaching is mitigated and we can probe each residue multiple times, thereby increasing the precision. After obtaining a sufficient number of FRET events, the FRET fingerprint can be constructed, reporting on the distance of each target amino acid from the reference point. Then the microfluidic chamber is washed and a new imaging solution is injected to probe a second amino acid (e.g. Lysine) (**Figure 1C**). The FRET X cycle can be repeated for any number and type of amino acids, as long as they are labeled with orthogonal DNA sequences. The detection of multiple types of amino acids improves the uniqueness of a protein fingerprint, thereby enhancing the chance of identification. The resolved FRET efficiencies for each amino acid are combined to generate a protein fingerprint, with which a protein can be identified against a reference database (**Figure 1D**).

**Figure 1.**
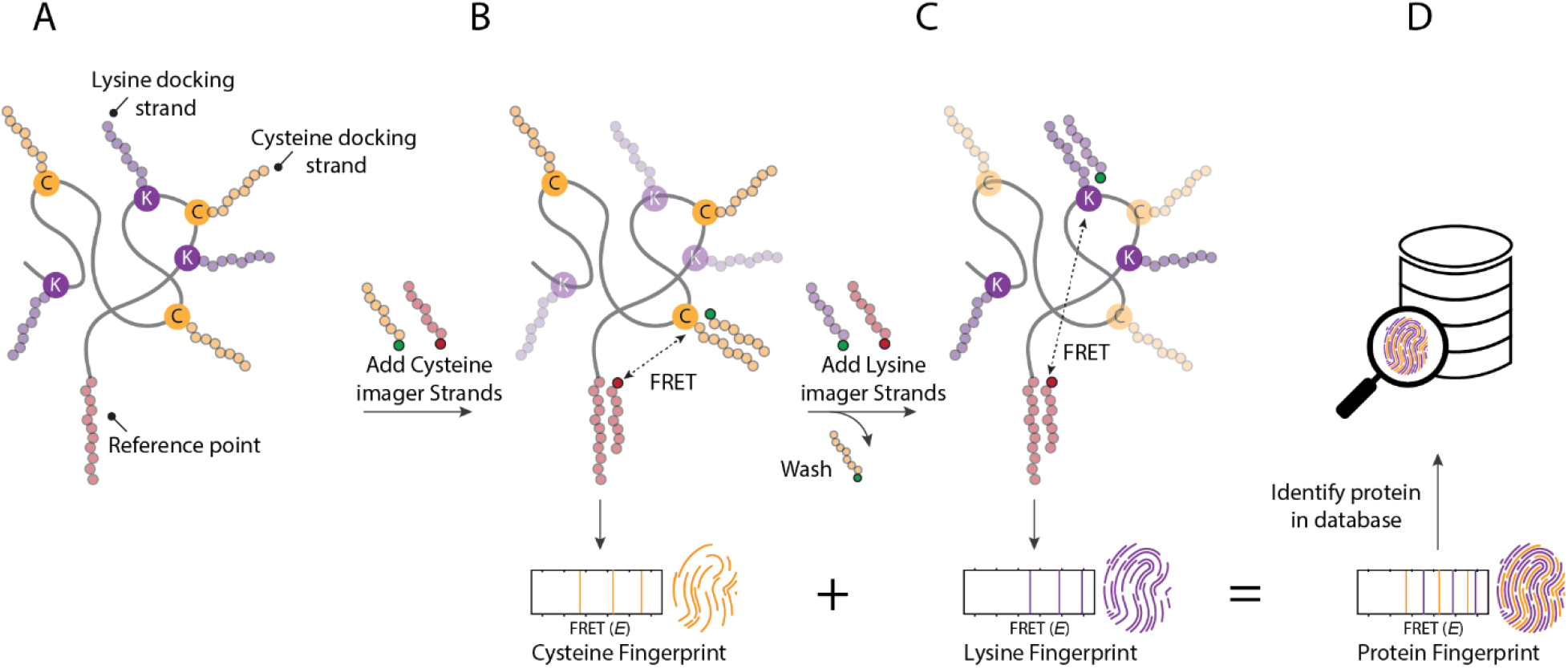
The concept of FRET X for protein fingerprinting. **(A)** A subset of amino acids (here cysteines and lysines) are labeled with orthogonal DNA sequences which function as docking sites for complementary, fluorescently labeled imager strands. Another orthogonal DNA sequence is conjugated to one of the protein termini, which serves as an acceptor docking site and facilitates immobilization of the protein to a microfluidic device. **(B)** In the first round of FRET X imaging, imager strands that hybridize with the cysteine docking site (yellow circles) and those that hybridize with the reference point (red circles) are injected in the microfluidic chamber. Both the donor and acceptor labeled imager strands transiently interact with their complementary docking strands. When both are present at the same time, FRET can occur and the FRET efficiency is determined between a cysteine and the reference point. Each of the three FRET pairs is separately probed, giving rise to a number of FRET efficiencies (*E*), which constitute the cysteine fingerprint. **(C)** The chamber is washed and FRET X imaging is repeated to probe the lysine fingerprint. This FRET X cycle can be repeated to probe additional amino acids and generate additional fingerprints. **(D)** The FRET efficiencies for individual amino acids are combined to produce a protein fingerprint that can be mapped against a reference database to identify the protein.

### Fingerprinting simulations

The usefulness of our method hinges on its ability to discern FRET X fingerprints derived from many different proteins. We run simulations to assess this scenario. Simulating the FRET X fingerprint for a given protein is a complex endeavor, as the fingerprint incorporates both sequence and structural information. While protein structure prediction has seen major advancements recently, cutting-edge methods^16,17^ remain too computationally costly to assess many proteins. Furthermore, they cannot account for the presence of conjugated DNA tags. Instead, we opted to use a computationally much less intensive lattice modelling approach^18^, in which each residue is represented as a single pseudo-atom, restricted in space to only occupy the vertices of a lattice (**Supplementary Figure 1**). Such structures can be efficiently energy-minimized by a Markov chain Monte Carlo process. Despite their simplicity, past investigations have shown that lattice models can reproduce native protein folding behavior.^19-23^

The attachment of DNA tags to selected residues, as required to accurately model our approach, has not previously been included in lattice models. The precise effect of DNA tags on protein structure is unclear. However, we find that implementation at the coarse granularity required by lattice models may be built on three basic assumptions: that tags prefer to reside on the exterior of the protein, require sufficient unoccupied space to avoid steric hindrance and repel each other if situated closely together. In the lattice models thus produced, FRET values can then be estimated from the simulated dye positions. To simulate the read-out of FRET efficiencies at a given resolution, we bin efficiencies using the resolution as bin width. As we have shown in previous work that a resolution of one FRET percentage point (0.01 E) is achievable, we set the resolution of fingerprints to 0.01 E in simulations, unless otherwise noted. As FRET X allows for orthogonal read-out of multiple residue types, the sampling can be repeated to produce the FRET fingerprints associated with different residue types. Analogously to experimentally obtained fingerprints, simulated FRET fingerprints for several residue types are then combined to serve as features for automated classification algorithms.

## Results

### Experimental FRET X fingerprinting of model peptides

To demonstrate the concept of protein fingerprinting using FRET X and to compare results with computational predictions, we designed an assay where DNA labeled peptides were immobilized on a PEGylated quartz surface via biotin-streptavidin conjugation (**Figure 2A**). Each peptide contains an N-terminal lysine for the attachment of a DNA-docking strand, to allow for the transient binding of an acceptor (Cy5)-labeled imager strand. Additionally, an orthogonal DNA-docking strand was conjugated to a cysteine residue in the peptide to facilitate transient binding of the donor (Cy3)-labeled imager strands (**Figure 2A**). The donor and acceptor imager strands were designed to exhibit a dwell time of ~ 2 s (**Supplementary Figure 2**), so that dyes could be frequently replenished. Furthermore, to increase the probability of the presence of the acceptor imager strand upon donor imager strand binding and allow for FRET detection, we injected 10-fold molar excess of the acceptor imager strand over the donor imager strand. Short-lived FRET events were recorded with single-molecule total internal reflection microscopy upon binding of both donor and acceptor labeled imager strands to the immobilized target peptide.

**Figure 2.**
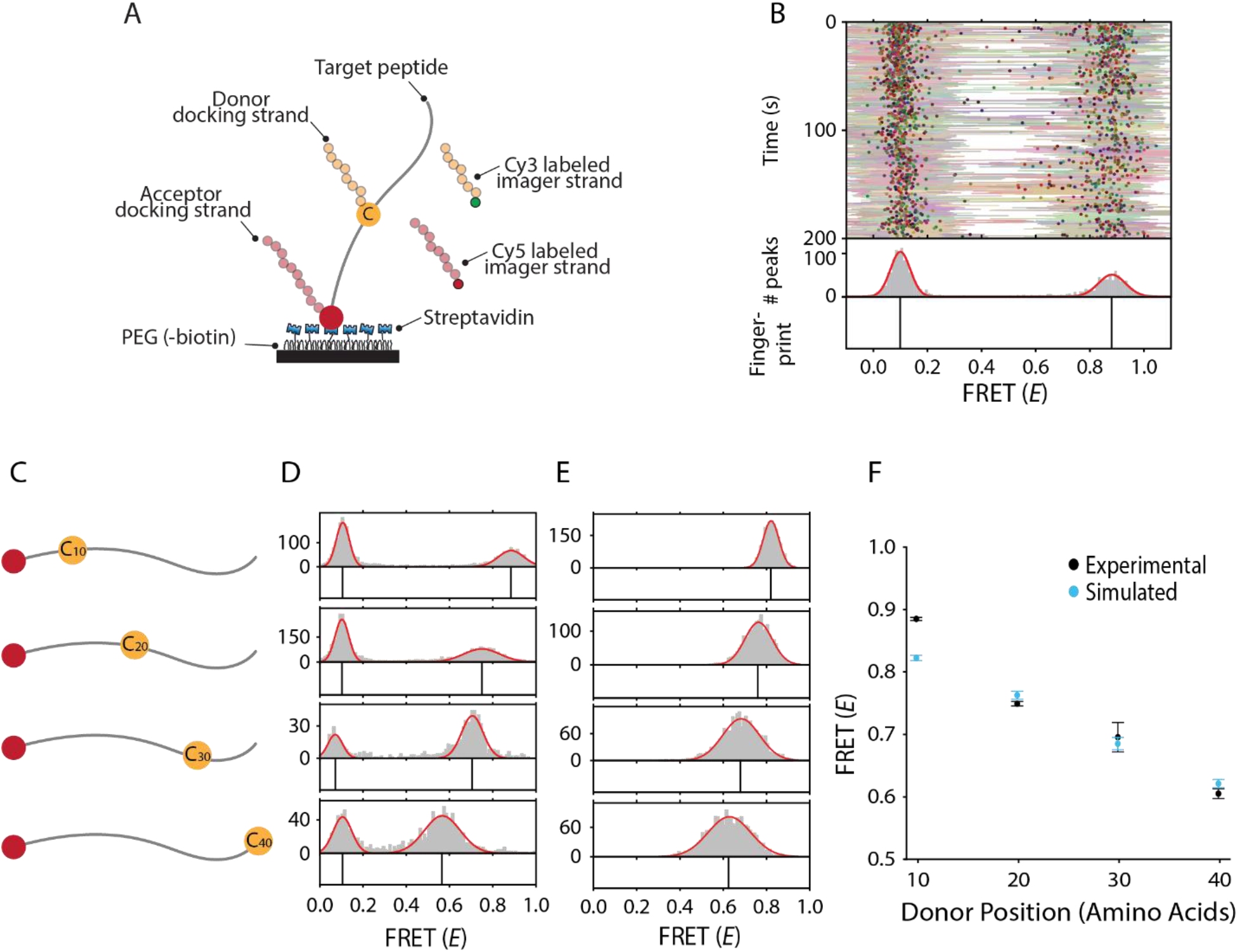
Model peptides can be fingerprinted with FRET X. **(A)** Depiction of the experimental system for peptide fingerprinting. The target peptide is immobilized through conjugation of its N-terminal biotin with the streptavidin on the PEGylated surface. The donor (Cy3) labeled imager strand (green) can bind to the DNA docking site on the cysteine, while the acceptor (Cy5) labeled imager strand (red) can hybridize to the docking site on the lysine. Simultaneous binding generates short FRET events and is observed with total internal reflection microscopy. **(B)** Representative kymograph for a peptide with a cysteine that is 10 amino acids separated from the acceptor binding site. The FRET efficiency for each data point in a binding event (lines) and the mean FRET efficiency from all data points in a binding event (dots) are indicated as a function of time. A Gaussian distribution (0.88 ± 0.05) is fitted on a histogram of average FRET efficiencies per FRET event. The means of the Gaussians are plotted in a separate panel (bottom) and are referred to as the FRET fingerprint of the peptide. The FRET population on the left is caused by donor leakage into the acceptor channel. **(C)** Our four model peptides have a lysine at the N-terminus and a cysteine at position 10, 20, 30 or 40. **(D)** Experimental distributions and fingerprints for each peptide show a downward trend in FRET (*E*) for increasing FRET pair separation (0.89 ± 0.06, 0.75 ± 0.08, 0.72 ± 0.03, 0.57 ± 0.08). **(E)** The simulated distributions and fingerprints for the four peptides show a similar downward trend. **(F)** Experimental and simulated data correlate well. Standard deviation of experimental data points is over four kymographs (each consisting of hundreds of events). Experiments were performed on separate days.

Next, we plotted a kymograph to visualize the FRET efficiency of each binding event in a target peptide (**Figure 2B**). The FRET efficiency for each data point (**Figure 2B**, lines) and the mean efficiency per binding event are calculated (**Figure 2B**, circles). A histogram of the mean FRET efficiency per binding event shows distinct FRET populations. Gaussian distributions were fit to resolve peak centers with high resolution^14^, which then constitute the fingerprint of the peptide (**Figure 2B**, bottom panel).

To demonstrate the ability of FRET X to distinguish different peptides with varying FRET pair separations, we designed four model peptides. These peptides had an incrementing distance, in steps of 10 amino acids, between donor and acceptor docking strands (**Figure 2C**). First, we performed singlemolecule experiments to obtain experimental FRET fingerprints and found a clearly discernible peak for each peptide (**Figure 2D and Supplementary Figure 3**). Then we simulated FRET fingerprints for the same sequences using our simulation pipeline and found a similar trend. We only fine-tuned the parameters for the repulsion effect between tags to minimize the difference with experimental values (**Figure 2E**). In both simulations and experiments we observe a monotonous decrease in FRET efficiency for increasing FRET pair separation. Furthermore, the experimentally obtained fingerprints generally correlate well with values found by simulations (**Figure 2F**). Since for each peptide the minimum inter-peptide difference in FRET (*E*) is larger than the maximum standard deviation, we find that we can distinguish these four peptides by their FRET fingerprint.

### Fingerprinting simulation of protein spliceoforms

We set out to evaluate the performance of our method for targeted proteomics, based on simulations. For this we sought to identify the different spliceoforms of the apoptosis regulator Bcl-2 (UniProt ID: Q07817), which are potential biomarkers for cancer^23^ and are likely to produce different fingerprints. While BCL-X_L_ is an anti-apoptotic regulator, both BCL-X_S_ and BCL-X_b_ are pro-apoptotic factors.^24,25^ The ratio between these factors is important for cell fate. We simulated simultaneous labeling of cysteine (C) and lysine (K) to create C+K fingerprints for each of the spliceoforms, BCL-X_L_, BCL-X_S_, and BCL-X_B_ (**Figure 3A and B**). As the spliceoforms differ in the numbers and locations of C and K residues, we expected their fingerprints to be dissimilar. This was indeed the case in simulation (**Figure 3C**). Fingerprints do vary across individual molecules of the same spliceoform; however, the fingerprints remain sufficiently characteristic to identify each spliceoform by eye (**Supplementary Figure 4A**). We also trained and tested a support vector machine (SVM) classifier on 10 replicates in a 10-fold cross validation scheme and attained an accuracy of 100%.

**Figure 3.**
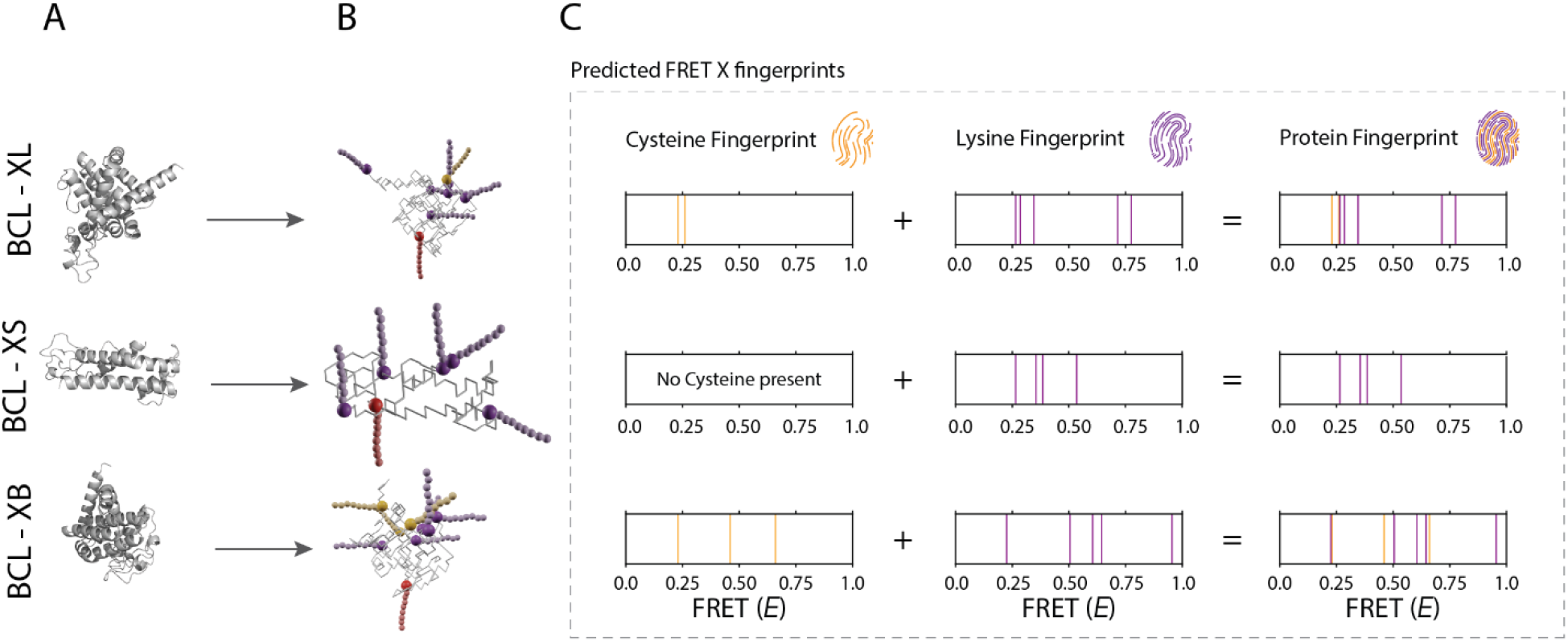
Representative FRET (*E*) fingerprints for three spliceoforms of BCL-X. **(A)** Fully atomic structure for BCL XL, Xs and Xb (from top to bottom) as predicted by the RaptorX structure prediction tool. **(B)** Energy-optimized lattice model structures with DNA-docking strands attached to cysteines (orange) and lysines (purple). The reference acceptor docking strand (red) is added to the N-terminus of the proteins. **(C)** The simulated fingerprint for spliceoform of the BCL proteins. Fingerprints are based on averaged donor-acceptor distances in 100 structural snapshots of Markov chain-generated lattice model structures.

We then simulated a more difficult scenario, in which we attempted to classify fingerprints for six spliceoforms of PTGS1 (UniProt ID: P23219).^26^ Although the higher number of C and K residues made discrimination of fingerprints by eye harder, an SVM trained and tested in a 10-fold cross validation scheme was still able to separate the six spliceoforms with 100% accuracy (**Supplementary Figure 4B**).

### Analysis of simulated protein mixtures

To evaluate a test case displaying a complexity closer to that found in a single cell, we selected all UniProt human proteome (ID: UP000005640) entries that were linked to a single-chain structure in the RCSB protein database and for which lattice modeling was able to find a configuration without steric hindrance of docking strands (n = 313). Based on available targeted residue labeling chemistries and relative residue frequencies in naturally occurring proteins, we simulated labeling schemes involving cysteine (C), lysine (K) and arginine (R). For each protein we generated fingerprints based on 10 separately simulated molecules, after which we trained and tested an SVM classifier in a 10-fold cross validation scheme. Here we report overall classifier accuracy. To identify the subset of proteins for which our method works well, we also report the number of well-identifiable proteins, i.e. those for which more than 5 of the replicates were identified correctly.

We find that our classifier performs at 45% accuracy on C-labeled proteins. Of 313 proteins, 126 were well-identifiable, indicating that labeling only C-residues is sufficient to consistently recognize this subset of proteins (**Figure 4A**, orange circle). 57 proteins did not contain C residues and are thus impossible to identify using C-labeling. The remaining 130 poorly identifiable proteins generally produced fingerprints containing few FRET values or highly variable fingerprints, the latter indicating a lack of structure stability.

**Figure 4.**
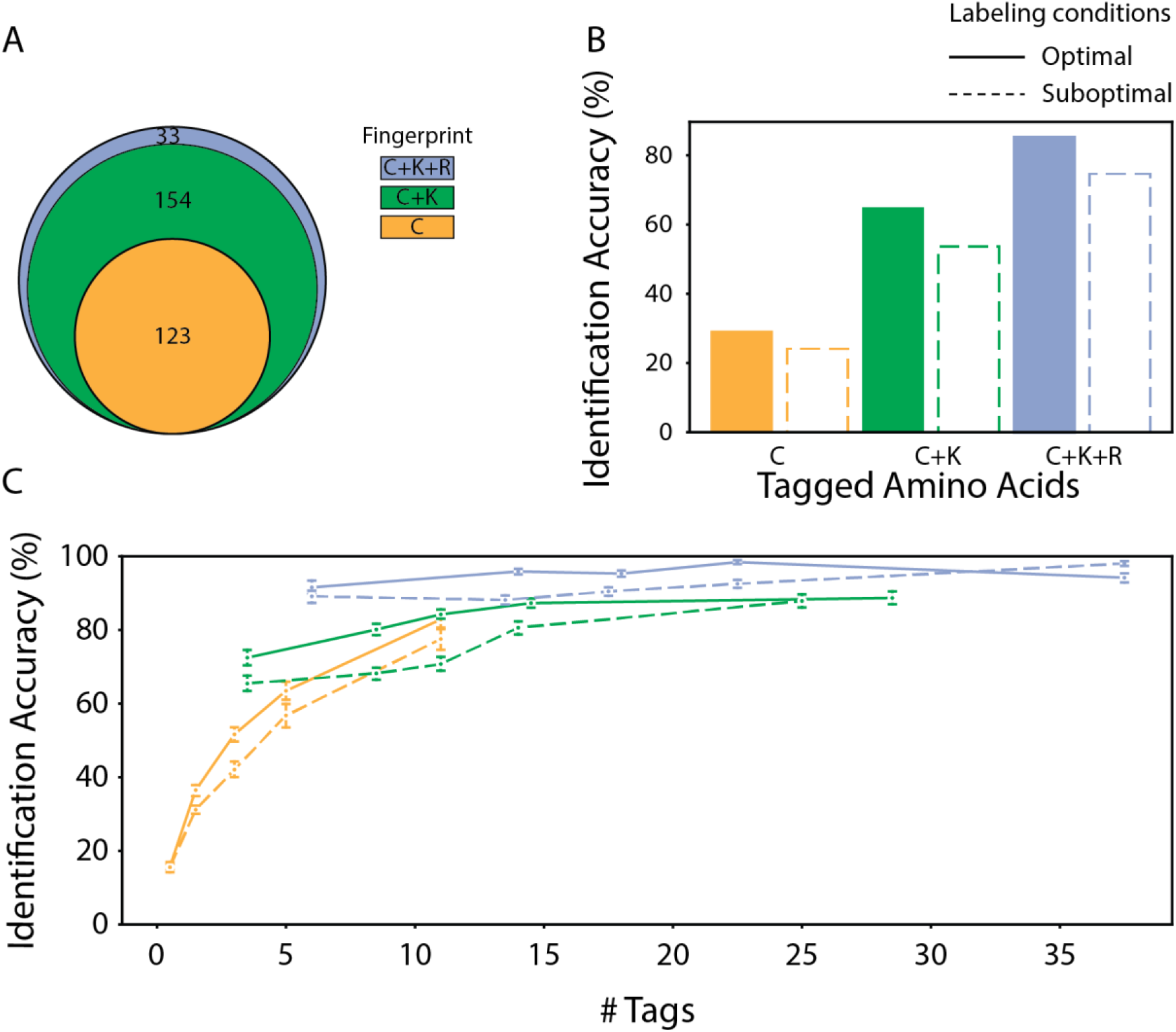
FRET X fingerprinting simulation results assuming optimal and suboptimal experimental conditions. FRET X fingerprint classifier cross-validation performance measures are shown for three combinations of tagged residue types - C, C+K, and C+ K+R - and two labeling qualities - “optimal”, where all targeted residues and no off-target residues were labeled, and “suboptimal”, where erroneous labeling occurred following the rules in Supplementary Table 3. **(A)** Venn diagram showing numbers of proteins that were found to be well-identifiable, i.e. that were correctly identified in more than 5 of 10 cross-validation folds. The total number of proteins is 313. **(B)** The identification accuracy of proteins under optimal and suboptimal labeling conditions. **(C)** Average classifier accuracy as a function of the number of tagged residues in structures, aggregated in five groups with similar numbers of tags. Whiskers denote two standard deviations.

When C+K or C+K+R residues were labeled, accuracy rose to 82% and 95% respectively (**Figure 4B**). As expected, fingerprints are more likely to obtain a characteristic signature if distances for more residue types are tracked. Numbers of well-identifiable fingerprints also rose to 278 and 312 out of 313 respectively. Regardless of which residue types are labeled, we find that proteins containing more tagged residues can be identified with higher accuracy **(Figure 4C)**.

### Robustness against suboptimal experimental conditions

To investigate the effect of labeling errors, we ran simulations for a suboptimal labeling scenario, with a 90% probability of labeling the target residue and a certain non-zero probability to label non-target residues (C: 1%, K:1%, R:0.5%, **Supplementary Table 3**). For C and K these probabilities were based on experimentally determined efficiencies and specificities found in literature.^27-29^

Overall, we find that labeling errors incur a modest decrease in classifier performance; for C, C+K and C+K+R labeling, accuracy drops from 45%, 82% and 95% to 39%, 74% and 91% respectively (Figure 4B). This indicates that FRET fingerprints - particularly those gained from C+K+R labeling - contain the redundant information required to mitigate the effect of imperfect labeling (Figure 4C). We also investigated the effect of decreased measurement resolution, however only after reducing resolution far beyond experimentally attainable levels - past 0.10 E - did we find severe reductions in accuracy (Supplementary figure 5).

## Discussion

Here we present a new protein fingerprinting approach that determines the location of amino acids within a protein structure using FRET X. We provide evidence of its ability to identify proteins in heterogeneous mixtures using simulations and demonstrate its technical feasibility by producing experimental fingerprints for designed peptides.

We experimentally demonstrate fingerprinting of peptides of 40 amino acids long and observe a monotonous decrease in FRET efficiency. This trend is supported by simulations and suggests that our model peptide has a relatively linear conformation. These peptides do not exhaust the lower end of the FRET-efficiency domain, which implies that larger peptides and proteins with increased FRET pair separation can be fingerprinted. While most proteins are considerably larger than 40 amino acids, they usually adopt a globular structure, which reduces the FRET pair separation. The average protein is estimated to have a diameter of 5 nm^30^, while the FRET dyes (Cy3-Cy5) used here are expected to be accurate at distances of up to ~7 nm.^1211^ Therefore, our FRET fingerprinting approach could be suitable for the identification of a large set of human proteins. This notion is substantiated by the simulations run using our lattice model, which shows that also for larger proteins the FRET fingerprints remain discernible.

We show that simulated fingerprints are sufficiently unique and reproducible to consistently identify the majority of the proteins in our simulation pool. Moreover, this result could be achieved by labeling up to three types of amino acids: cysteine, lysine and arginine, all of which can be targeted for specific labeling using existing chemistries.^4,27^ Interestingly, even if only cysteine is labeled we find that a subset of proteins remained consistently identifiable, although labeling additional residue types does increase accuracy, the number of identifiable proteins and robustness against labeling errors. It should also be noted that the set of residue types targeted for FRET X fingerprinting can be expanded even further; labeling of e.g. methionine^31^ or tyrosine^32^ may be employed to further increase accuracy or tailor our method to the detection of a given target protein. For our simulations we investigated proteins for which the structure had already been determined; however, in our experimental system, a microfluidic chamber with non-physiological conditions, proteins may adopt a different structure or a set of several different structures, creating a discrepancy between simulated and experimental fingerprints. However, it is primarily the uniqueness and reproducibility of a fingerprint that is important for protein identification, not necessarily its predictability from a known structure. Furthermore, we expect that as the diversity of a sample decreases from several hundreds to tens of different proteins through sample fractionation, the fingerprint uniqueness and thereby the fraction of correctly identified proteins sharply increases. Adequate sample preparation and purification to reduce sample complexity will therefore be important for more targeted approaches.

A far-reaching goal of the proteomic community is to detect and analyze all proteoforms that can be derived from a single protein encoding gene.^2^ Most proteoforms have subtle differences, e.g. alternative splicing or post translational modification, and are difficult to detect with current technologies, such as ELISA, MS or native MS.^33^ We have shown that FRET X has the ability to distinguish peptides based on the location of a single cysteine, a subtlety akin to those found in many isoforms, and we have shown two cases in which clinically relevant spliceoforms are well distinguishable based on their simulated FRET X fingerprints. This suggests that our FRET X fingerprinting platform would be a suitable complementary technique for the detection of clinically relevant proteoforms.

## Acknowledgements

We thank Sung Hyun Kim for fruitful discussions and feedback. C.J. and D.R. acknowledge funding from NWO-I (SMPS).

