## Supplementary information for "Evaluation of FRET X for Single-Molecule Protein Fingerprinting"

| Supplementary Methods | |
|---|---|
| Figure S-1 | **Pseudo-atoms on a cubic lattice (A) and a body-centered cubic lattice (B).** |
| Figure S-2 | **Single-molecule binding kinetics of FRET X imager strands.** |
| Figure S-3 | **Representative kymographs of individual peptides.** |
| Figure S-4 | **Simulated FRET X fingerprints for spliceoforms of BCL and PTGS1.** |
| Figure S-5 | **SVM classifier accuracy on simulated fingerprints for 313 proteins at different resolutions.** |
| Figure S-6 | **Schematic of FRET X fingerprinting simulation and classification pipeline used in this work.** |
| Figure S-7 | **Illustration of the tag repulsion implementation of the lattice model.** |
| Table S-1 | **Single-molecule peptide constructs.** |
| Table S-2 | **Single-molecule DNA constructs.** |
| Table S-3 | **Labeling probabilities under suboptimal conditions.** |

### Materials and Methods

#### Peptide Labeling

Custom designed polypeptides were obtained from Biomatik (Canada) and had a constant backbone sequence (see Supplementary Table 1), differing only in the cysteine substitutions. Cysteine residues of the polypeptides were reduced with 40-fold molar excess Tris(2-carboethyl)phosphine (TCEP) for 30 minutes and then donor-labeled with 6-fold molar excess monoreactive maleimide-(5') functionalized DNA in 50 mM HEPES pH 6.9 overnight at room temperature. The acceptor docking strand was labeled onto a single lysine that is located at the N-terminus of the peptide. For this, Dimethyl sulfoxide (DMSO) was added to 50 % (v/v) and the pH was increased to pH 7.5 through the addition of NaOH. Next, we added monoreactive N-Hydroxysuccinimide (NHS)-ester functionalized Dibenzocyclooctyne (DBCO) (Sigma Aldrich, Germany) in a 25-fold molar excess and incubated for 6 hours at room temperature. Free NHS-DBCO was removed by using C18 bed micropipet tips (Pierce) according to manufacturer's protocol. Finally, monoreactive Azidobenzoate-(5') functionalized-DNA was added in 5-fold molar excess and incubated overnight at room temperature. See Supplementary Table 1 and 2 for the full list of substrates.

#### Single-Molecule Setup

All experiments were performed on a custom-built microscope setup. An inverted microscope (IX73, Olympus) with prism-based total internal reflection was used. In combination with a 532 nm diode-pumped solid-state laser (Compass 215M/50mW, Coherent). A 60x water immersion objective (UPLSAPO60XW, Olympus) was used for the collection of photons from the Cy3 and Cy5 dyes on the surface, after which a 532 nm long pass filter (LDP01-532RU-25, Semrock) blocks the excitation light. A dichroic mirror (635 dcxr, Chroma) separates the fluorescence signal which is then projected onto an EM-CCD camera (iXon Ultra, DU-897U-CS0-#BV, Andor Technology). A series of EM-CDD images was recorded using a custom-made program in Visual C++ (Microsoft).

#### Single-Molecule Data Acquisition

Single-molecule flow cells were prepared as previously described.[34,35] In brief, to avoid non-specific binding, quartz slides (G. Finkerbeiner Inc) were acidic piranha etched and passivated twice with polyethylene glycol (PEG). The first round of PEGylation was performed with mPEG-SVA (Laysan Bio) and PEG-biotin (Laysan Bio), followed by a second round of PEGylation with $MS(PEG)_4$ (ThermoFisher). After assembly of a microfluidic chamber, the slides were incubated with 20 μL of 0.1 mg/mL streptavidin (Thermofisher) for 2 minutes. Excess streptavidin was removed with 100 μL T50 (50mM Tris-HCl, pH 8.0, 50 mM NaCl). Next, 50 μL of 75 pM DNA-labeled peptide was added to the microfluidic chamber. After 2 minutes of incubation, unbound peptide and excess Azide-DNA from the earlier click reaction was washed away with 200 μL T50. Then, 50 μL of 10 nM donor labeled imager strands and 100 nM acceptor labeled imager strands in imaging buffer (50 mM Tris-HCl, pH 8.0, 500 mM NaCl, 0.8 % glucose, 0.5 mg/mL glucose oxidase (Sigma), 85 ug/mL catalase (Merck) and 1 mM Trolox (Sigma)) was injected. All single-molecule FRET experiments were performed at room temperature (23 ± 2 °C).

**Data analysis**

Fluorescence signals are collected at 0.1-s exposure time unless otherwise specified. Time traces were subsequently extracted through IDL software using a custom script. Through a mapping file, the script collects the individual intensity hotspots in the acceptor channel and pairs them with intensity hotspots in the donor channel, after which the time traces are extracted. During the acquisition of the movie, the green laser is used to excite the Cy3 donor fluorophores. For automated detection of individual fluorescence imager strand binding events, we used a custom Python code (Python 3.7, Python Software Foundation, https://www.python.org) utilizing a two-state K-means clustering algorithm on the sum of the donor and acceptor fluorescence intensities of individual molecules to identify the frames with high intensities.[31] To avoid false positive detections, only binding events that lasted for more than three consecutive frames were selected for further analysis. FRET efficiencies for each imager strand binding event were calculated and used to build the FRET kymograph and histogram. Populations in the FRET histogram are automatically classified by Gaussian mixture modeling.

**Simulations**

Fingerprinting simulations were generated using a lattice folding model written in Python 3.7. Simulation and analysis code are freely available at https://github.com/cvdelannoy/FRET_X_fingerprinting_simulation. A protein folding simulation was implemented to incorporate DNA-tags attached to certain residues and account for their effect on the protein structure. Lattice models were used because of the far lower computational power needed for folding simulations compared to fully atomistic models allowing unrestricted movement, which is attained by reducing each amino acid to a pseudo-atom and restricting its possible positions to the vertices of a lattice. Such models have previously been used in applications where low computational requirements were essential.[16–20] The procedure starts with a fully atomistic native structure, which is converted to a lattice structure with tagged residues marked. This structure is then refolded by making local modifications and calculating the effect these have on the model energy ($E_{tot}$), as calculated by an energy function. Modifications that decrease $E_{tot}$ are accepted, whereas those that increase $E_{tot}$ are more likely to be discarded the more they increase $E_{tot}$. The procedure ends when all DNA-tags fit in the structure without causing steric hindrance. Aspects of the modeling procedure are described in more detail below.

**Lattice structure**

The lattice modeling procedure employed here largely resembles those in previously published applications.[19] In particular, the model developed by Abeln et al.[19] was used as a starting point, however the cubic lattice was replaced by a novel body-centered cubic (BCC) lattice (**Supplementary Figure 6**). The octahedral unit cell of a BCC lattice borders eight neighboring cells through its hexagonal faces and four through its square faces. However, only connections through hexagonal faces are considered, as this allows all bonds to be of the same length. As a result, only even coordinates in the lattice are valid vertices for residue placement.[33] This implementation increases the number of contacts that each non-endpoint residue can make from four to six (not including immediately neighboring residues) and increases the number of directions into which a bond may extend. The resulting increased flexibility allows lattice models to more closely resemble native folds. Moreover, alpha helices are represented better as the BCC lattice allows structures that make one regular turn per five residues.

##### Tag implementation

As the precise effect of the presence of DNA-tags on protein structure is unclear, we relied on several basic assumptions to include them in the model. First, we assume that DNA-tags prefer to reside in the periphery of a protein due to their polar backbones. Thus, labeling an internal residue should alter local structure to accommodate sufficient space from the residue to the surface, while tagging a residue that already resides on the protein surface should affect the structure less severely. This was implemented by adding a substantial energy penalty if a tagged residue did not have space for a DNA tag to reach the periphery of the structure without clashing with the main chain. Secondly, we assume that tags will electrostatically repel each other. This is represented by introducing a minimum angle and dihedral between tag pairs that are spatially close together in a given configuration (**Supplementary Figure 7**). To parameterize this effect, we compared predicted fingerprints of 40-residue model peptides to the presented experimental data and found that values are reproduced well if at least a 70° angle and dihedral are enforced between tags situated within 20Å of each other.

##### Simulated labeling scenarios

Two labeling scenarios are employed in this work. Under the optimal scenario, all target residues are labeled and no off-target labeling takes place. Under the suboptimal scenario, both labeling efficiency and specificity are decreased, following a similar procedure to Ohayon et al.[7]; each target residue has a 90% chance of being labeled by its dedicated chemistry, while some off-target labeling probability is defined for one or more other residue types. Where possible, efficiency and specificity parameters are based on literature (**Supplementary Table 3**).

##### Structure collection

We base the lattice models used in our fingerprinting simulations on fully atomistic structures as stored in the RCSB PDB. To obtain a dataset of relevant structures, we analysed all available PDB entries corresponding to entries in the Uniprot human proteome set (UP000005640). Of the 20,381 entries in the proteome, 7,133 solved structures were found. We further filtered this list on structure quality, retaining only those with an R-free value below 0.21, and removed structures with non-canonical residues as our model contains no energy modifiers for these residues. Lastly, quaternary structure is expected to be lost during sample preparation, thus to avoid having to model the effect of losing other chains on the tertiary structure of the target chain, we removed structures which were crystalized as a complex of multiple chains. After these filtering steps, 746 structures remained for our simulations.

A lattice models is derived from a fully atomistic structure by reducing it to its Cα positions and placing each Cα-atom on the nearest lattice vertex, while remaining connected to its neighboring Cα-atom, starting from the residue with the lowest index. Alpha helices are forced to remain intact on the lattice, by first translating involved Cα-atoms to a lattice-compliant helix and then minimizing the distance between their respective lattice positions simultaneously.

As no PDB structures are available for the 40-residue model peptides labeled in practical experiments, starting structures for these peptides were stretched configurations. Starting structures for BCL-X and PTGS1 spliceoforms were generated using the RaptorX structure prediction server.[36]

**Folding simulation**

After initialization of the lattice model, a Markov Chain Monte Carlo (MCMC) procedure is employed to minimize the structure energy $E_{tot}$.

$$E_{tot} = E_{AA} + E_{sol} + E_{ss} + E_{tag} + E_{reg}$$

Residue interaction and residue-solvent interaction terms $E_{AA}$ and $E_{sol}$ are summed pairwise interaction terms between contacting residues or residue-solvent contacts, the magnitudes of which are obtained empirically.[37] The secondary structure formation energy term $E_{ss}$ is adapted from Abeln et al.[19] and incurs an arbitrarily high energy bonus of -25 if an alpha helix or beta sheet is formed, but only if a given residue also was part of such a secondary structure in the native fold. An alpha helical residue incurs this bonus if the exact shape of the helix is formed (i.e. residue $i$ up to $i$+4 take the same relative orientation at each step), while a bonus for beta sheet formation is applied if non-neighboring beta-sheet residues are adjacent to each other. The tag energy term $E_{tag}$ incurs an arbitrarily high energy penalty of 100 for each residue impeding the shortest route from a tagged residue to the periphery of the structure. Lastly, the regularization term $E_{reg}$ incurs a penalty for large structural reorganizations occurring in a single MCMC step, as we found that this helps to retain the native fold as much as possible.

**Fingerprint extraction**

To account for the fact that a structure may adopt several conformations over the course of measurements, fingerprints are based on a series of structure snapshots. After the folding simulation has finished and the structure which accommodates all DNA-tags without steric hindrance is found, another 1,000 MCMC steps are performed. During these steps, snapshots are taken at intervals of 10 steps, thus measuring 100 slightly different conformations. For each snapshot, dye positions are chosen randomly from all accessible lattice directions. If tags are found to be closer than 20Å to each other, a minimum angle and dihedral angle of 70 degrees each between those tags is enforced (**Supplementary figure 7**). Distances between donor and acceptor dye positions are estimated from the snapshots and averaged, after which the FRET efficiency is calculated as follows:

$$E_{FRET} = \frac{1}{1 + (R/R_0)^6}$$

Here $R$ is the modeled distance between donor and acceptor dye and $R_0$ is the Förster radius, which characterizes the used FRET dye pair ($R_0$ assumed constant at 54Å for the Cy3-Cy5 FRET pair[12]). Finally, all FRET values are binned and normalized over the number of snapshots to produce the final fingerprint. The bin width is used here to represent the observation resolution. Resolution is fixed at 0.01 unless otherwise noted, as previous work has shown that such a resolution can be achieved using FRET X.[15] If multiple residue types are tagged, each residue type generates its own fingerprint which is binned separately.

**Classification**

To classify simulated fingerprints a support vector machine (SVM) was implemented using the scikit-learn package (v0.23.2.)[36]. In a ten-fold cross validation procedure, the SVM was fitted to a training set consisting of 90% of produced fingerprints and tested on a held-out test set. As a higher resolution is also more sensitive to noise by unstable fingerprints, the resolution is tuned during training in steps of 0.01 E to produce the highest training accuracy. To evaluate classifier performance, we calculated test accuracy, i.e. the number of correct classifications over total number of test examples. As this measure obscures whether classification mistakes are consistently made for certain proteins or are randomly distributed, we also determined which proteins were correctly classified in more than half of replicates, which we denote as well-identifiable proteins.

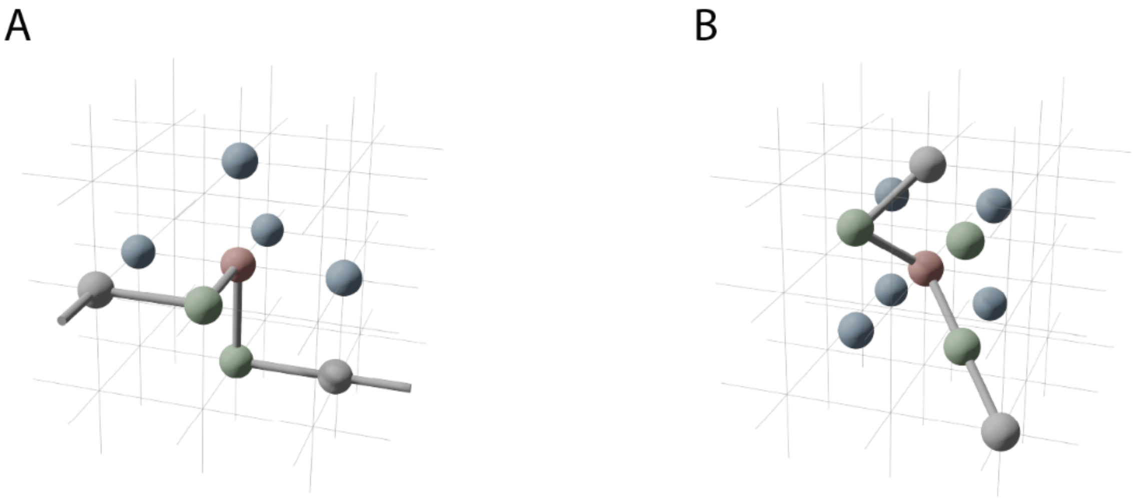


**Supplementary Figure 1: Pseudo-atoms on a cubic lattice (A) and a body-centered cubic lattice (B).** Shown are one main pseudo-atom (red) and its direct neighbors (green). For the main pseudo-atom, all possible adjacent pseudo-atoms (green) are depicted. Figures were generated in Blender 2.93.0.

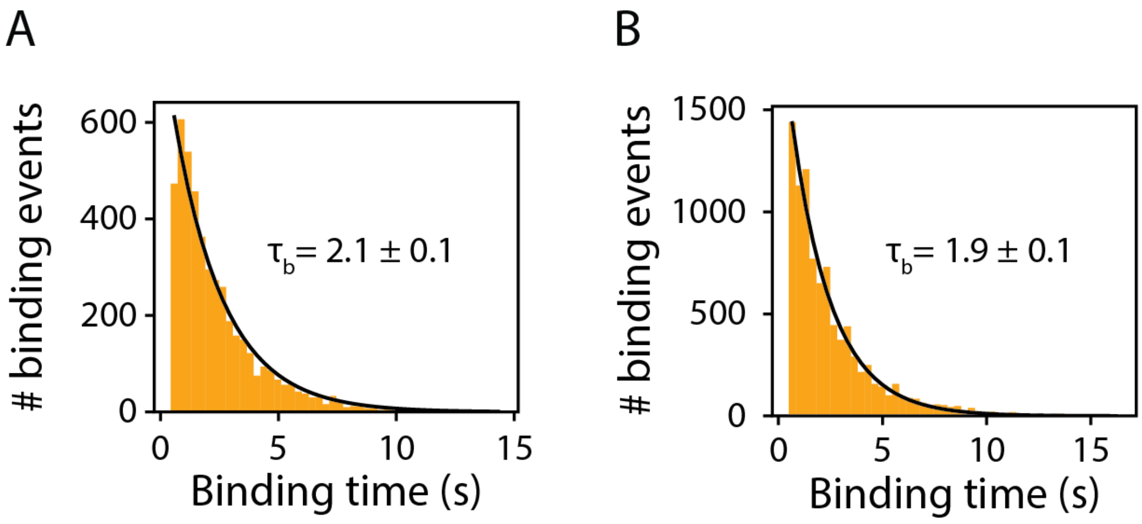


**Supplementary Figure 2: Single-molecule binding kinetics of FRET X imager strands.**
**(A)** Dwell-time histogram for the FRET X donor imager strand **(Supplementary Table 2)** fitted with a maximum likelihood estimation for a single exponential distribution (black line). Average ± standard deviation of four different estimations gives: 2.14 ± 0.07 s. The number of datapoints for this distribution: n = 4687 and peptide used was K1C40. **(B)** Dwell-time histogram for the FRET X acceptor imager strand **(Supplementary Table 2)** fitted with a maximum likelihood estimation for a single exponential distribution (black line). Average ± standard deviation of four different estimations gives: 1.92 ± 0.02 s. The number of datapoints for this distribution: n = 9477 and peptide used was K1C40.

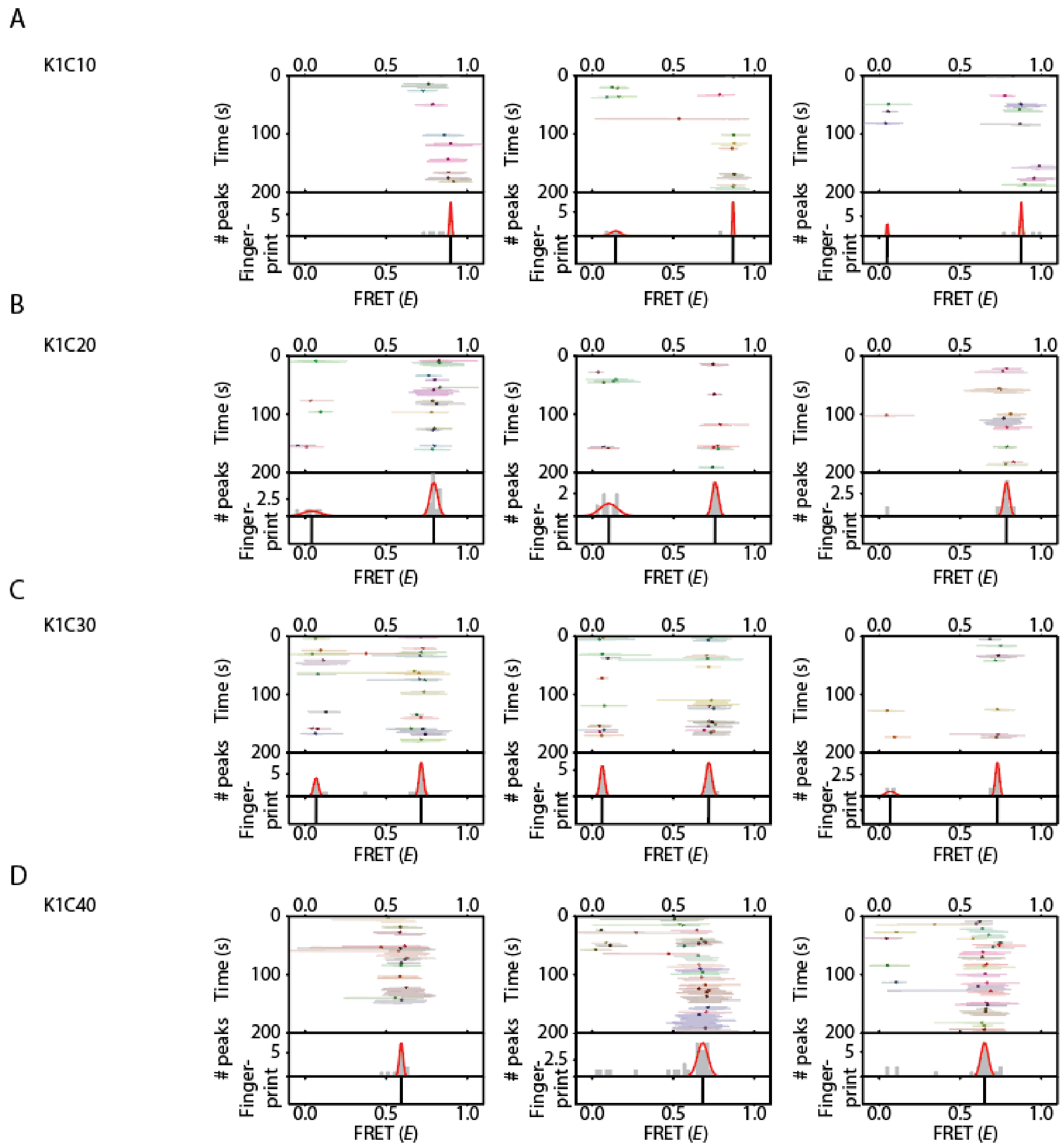


**Supplementary Figure 3: Representative kymographs of individual peptides.**

**(A-D)** Representative single-molecule FRET kymographs for each of the four peptides. The downward FRET (*E*) trend remains at the single-molecule level and the distribution of each individual molecule can be fitted with high precision (s.d. ≤ 0.03 for each distribution). The ensemble of many identical single-molecules results in the FRET-fingerprint (**Figure 2B**).

A

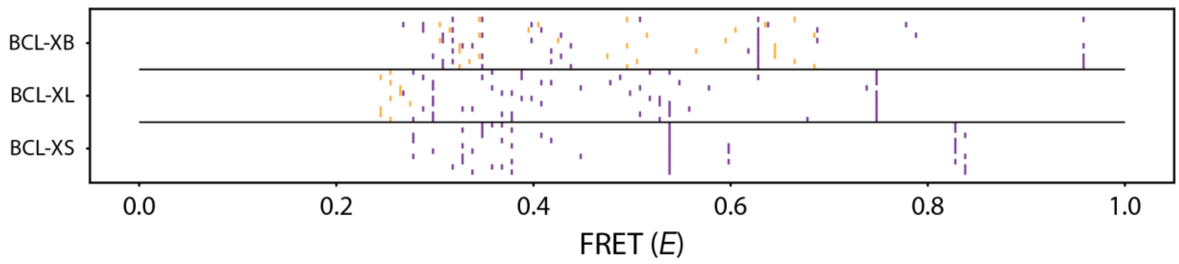


B

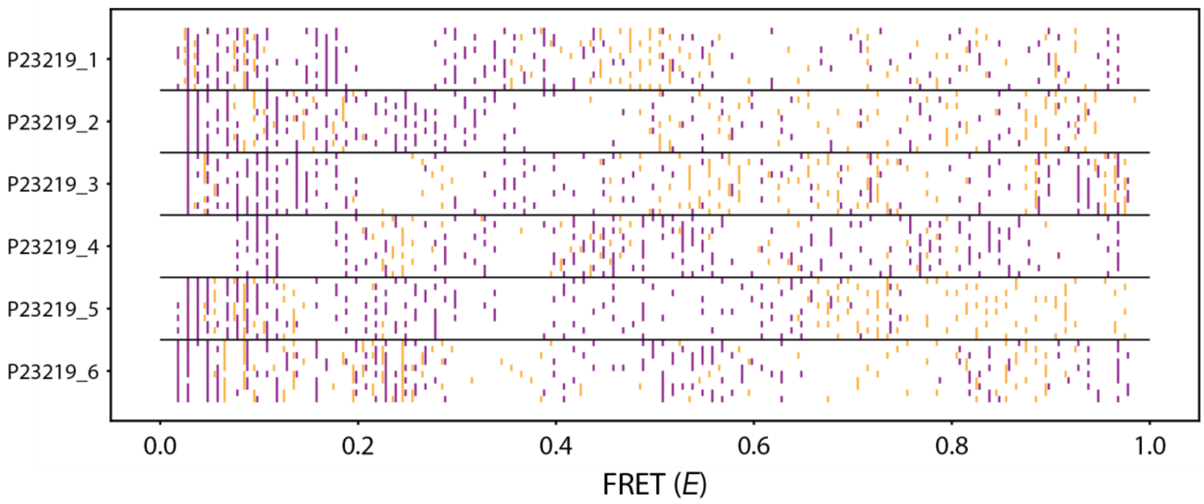


**Supplementary Figure 4: Simulated FRET X fingerprints for spliceoforms of BCL and PTGS1.**

**(A)** FRET X fingerprints for ten simulated molecules, one per horizontal line, of three BCL-X spliceoforms: BCL-$X_L$, BCL-$X_S$ and BCL-$X_B$. Cysteine and lysine-derived values are colored orange and purple respectively**. (B)** FRET X fingerprints for ten simulated molecules of six PTGS1 spliceoforms.

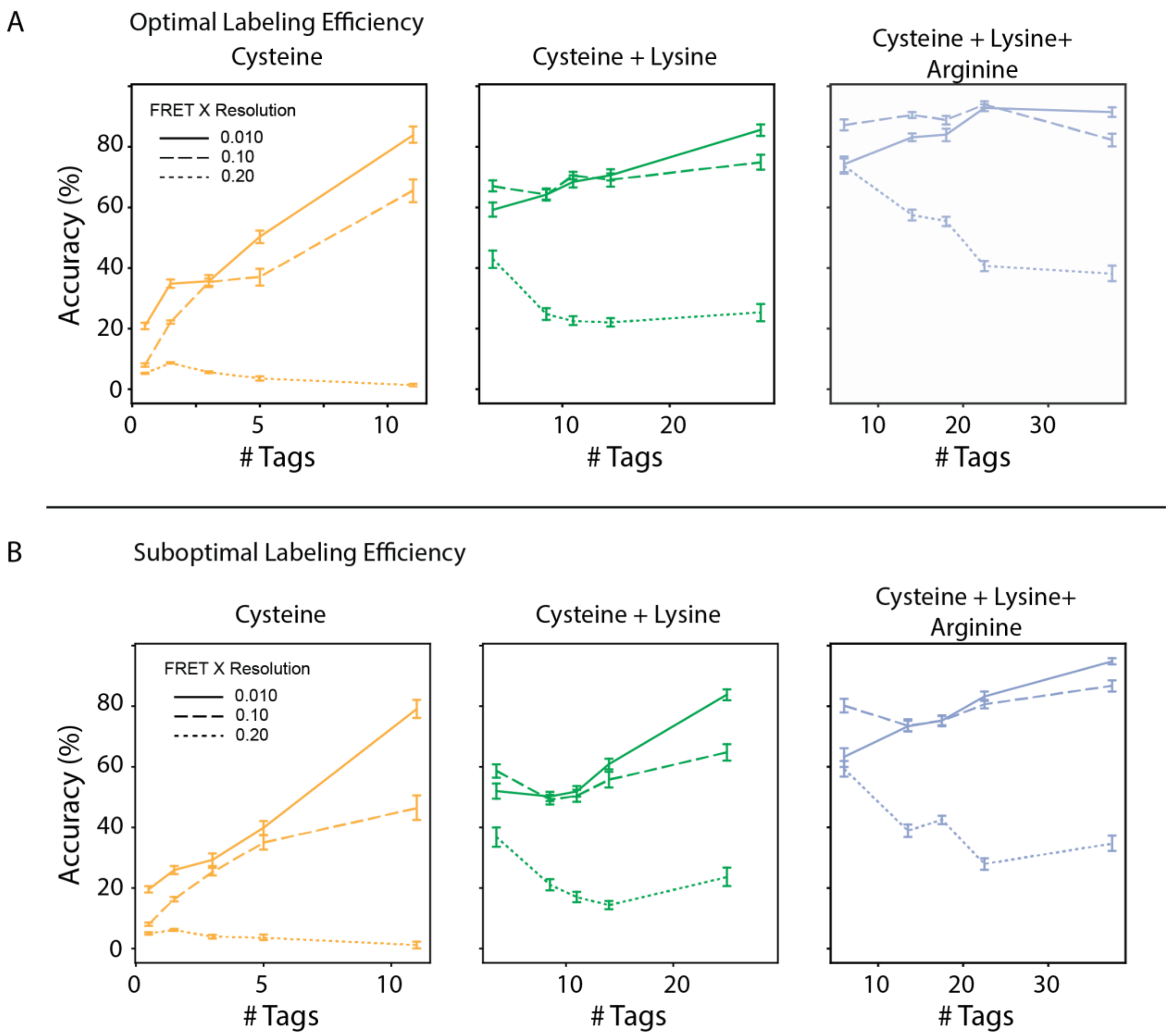


**Supplementary Figure 5: SVM classifier accuracy on simulated fingerprints for 313 proteins at different resolutions.** Average classifier accuracy versus the number of tagged residues in structures, aggregated in five groups with similar numbers of tags, at different resolutions. Data are shown for **(A)** optimal labeling quality (i.e. 100% efficiency, 100% specificity) and **(B)** suboptimal labeling quality (see supplementary table 3 for efficiency and specificity), for three combinations of tagged residues (C, C + K, and C+ K + R). Whiskers denote two standard deviations.

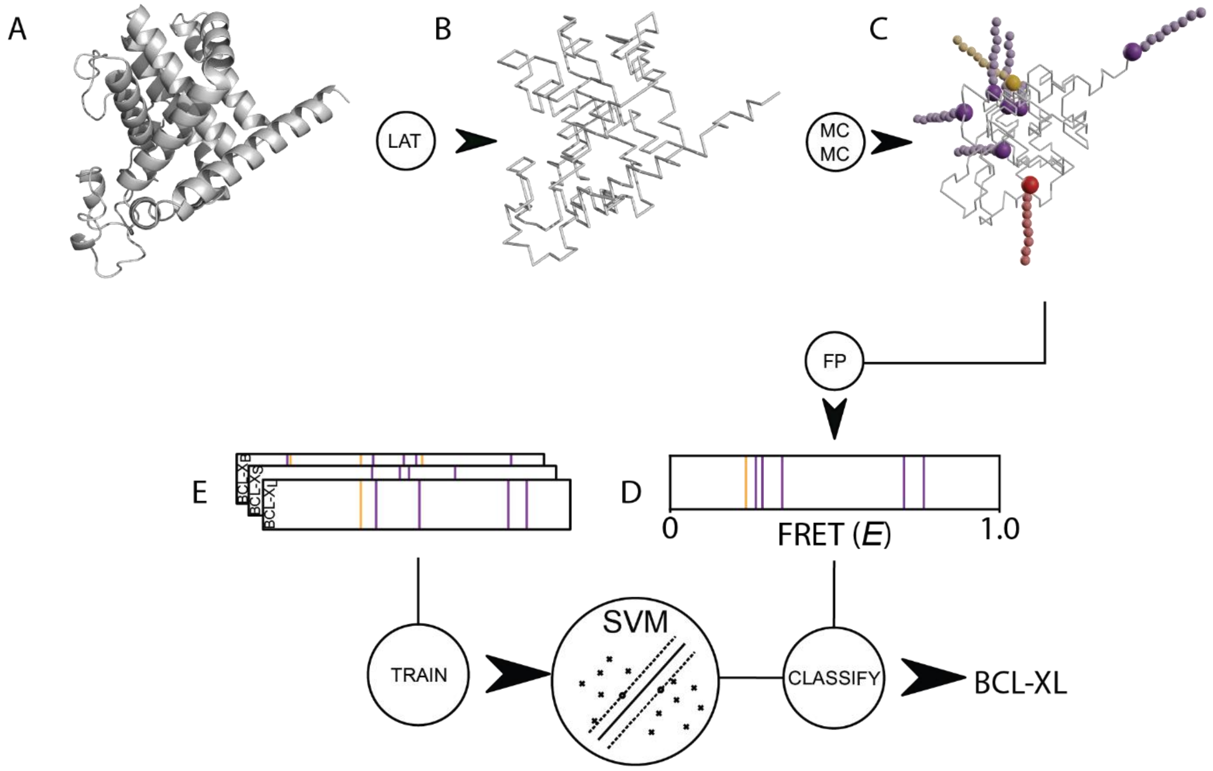


**Supplementary Figure 6: Schematic of FRET X fingerprinting simulation and classification pipeline used in this work**. **(A)** Simulation starts from a fully atomistic structure, **(B)** which is first converted into a lattice model. In the lattice model all residues are reduced to their Cα positions. **(C)** Residues to which docking strands must be attached are marked, after which the structure is randomly mutated using a Markov chain Monte Carlo (MCMC) process, until docking strands no longer experience steric hindrance from the rest of the structure. **(D)** The MCMC process then continues while snapshots are taken at regular intervals. Donor-acceptor dye distance for each dye pair is averaged over all snapshots and translated into a FRET efficiency. Combined, the FRET efficiencies form the final fingerprint for this molecule. **(E)** A support vector machine (SVM) is trained on a set of fingerprints with known identities, after which it can be used to classify unseen fingerprints.

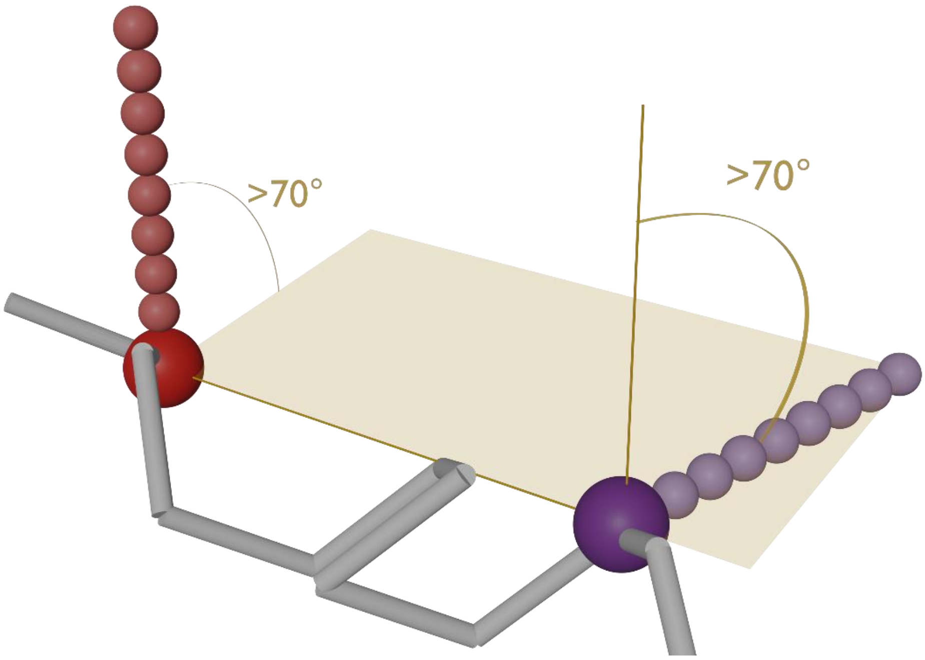


**Supplementary Figure 7: Illustration of the tag repulsion implementation of the lattice model.** If the distance between two tagged pseudo-atoms is found to be less than 20Å, both the angle and the dihedral angle should be larger than 70° to obtain a valid tag position. Figure was generated in Blender 2.93.0.

**Supplementary Table 1: Single-molecule peptide constructs.**

| Peptide | Sequence (N to C) | Modification | Supplier |
|---|---|---|---|
| K1C10 | KAGERDNFACHMALVPVAANDENYALAAAANDENYALAAA | Biotin-Ahx N-terminus | Biomatik (CAN) |
| K1C20 | KAGERDNFAPHMALVPVAACDENYALAAAANDENYALAAA | Biotin-Ahx N-terminus | Biomatik (CAN) |
| K1C30 | KAGERDNFAPHMALVPVAANDENYALAAACNDENYALAAA | Biotin-Ahx N-terminus | Biomatik (CAN) |
| K1C40 | KAGERDNFAPHMALVPVAANDENYALAAAANDENYALAAC | Biotin-Ahx N-terminus | Biomatik (CAN) |

**Supplementary Table 2: Single-molecule DNA constructs.**

| DNA Strand | Sequence (5' - 3') | Modification | Supplier |
|---|---|---|---|
| Donor imager strand | AGATGTAT | 3' Cy3 | Ella Biotech (GmbH) |
| Acceptor imager strand | AATGAAGA | 3' Cy5 | Ella Biotech (GmbH) |
| Donor docking sequence | TATACATCTAT | 5' Maleimide | Biomers.net (GmbH) |
| Acceptor docking sequence | TTCTTCATTACT | 5' Azidobenzoate | Biomers.net (GmbH) |

**Supplementary Table 3: Labeling probabilities under suboptimal conditions.**

| Chemistry | Target | P(target labeling) | Off-target residue | P(Off-target labeling) | Reference |
|---|---|---|---|---|---|
| Maleimide thiol reaction | C | 90% | K | 1% | Boutureira et al. [27] |
| NHS ester-mediated derivatization | K | 90% | S,Y,T | 1% | Abello et al.[28] |
| Arginine derivatization | R | 90% | Any | 0.5% | Thompson et al. [29] |
